# Curvature tuning in areas V2 and V4 of the developing macaque

**DOI:** 10.64898/2026.06.16.732692

**Authors:** A. Ezra Sutter, Gerick M. Lee, Timothy D. Oleskiw, Najib J. Majaj, Lynne Kiorpes, J. Anthony Movshon

**Author notes:** Equal contribution (A. Ezra Sutter, Gerick M. Lee). Department of Neuroscience, University of California, Berkeley, Berkeley, CA, United States. Department of Computer Science, University of Regina, Regina, SK, Canada.

## Abstract

Visual areas V2 and V4 are critical for the perception of visual forms in primates. Neurons in area V4 of macaque monkeys are often sensitive to the curvature of specific boundary segments within shapes, but it is unknown how curvature tuning is represented in the developing brain. To address this, we recorded multiunit neural activity from areas V2 and V4 of two macaque monkeys, at both 30 and 58 weeks of age, in response to shape stimuli which primarily varied in curvature along a single segment. Observable curvature tuning was adult-like from 30 weeks of age in both V2 and V4. We compared the tuning of sites to shapes presented at multiple positions. We saw evidence of position-invariant tuning in V4, but not in V2. Position invariance in V4 was stable from 30 weeks of age. Finally, we fit two models – a stimulus-centric model of boundary curvature tuning, and a simple linear model based on the spike-triggered average response to all stimuli. We found many sites in both V2 and V4 whose responses could be captured by one or both models, but no evidence of age-related changes in curvature tuning within the space of either model. Our results suggest that the neural representation of curvature in both areas reaches maturity soon after birth, and that object-centric representations of curvature first emerge in V4, not V2.

## INTRODUCTION

The primate visual system develops during early life (Dekker et al., 2020; Kiorpes, 2016). Studies in both humans (Dekker et al., 2020; Huber et al., 2023; Shimojo and Held, 1987; Zanker et al., 1992) and macaque monkeys (El-Shamayleh et al., 2010; Kiorpes, 2016; Kiorpes and Bassin, 2003; Kiorpes et al., 2012; Lee et al., 2024; Rodríguez Deliz et al., 2024) have repeatedly demonstrated that behavioral sensitivities across a wide variety of spatial vision tasks continue to mature during early life. In the macaque, these changes typically continue through the first year of life or longer. Conversely, physiological studies of development in the macaque visual system have found limited evidence of maturation after the first 8-16 weeks (Kiorpes and Movshon, 2004; Lee et al., 2024; Maruko et al., 2008; Movshon et al., 2005; Rodman et al., 1993; Rodríguez Deliz et al., 2025; Wang et al., 2019; Zhang et al., 2013, 2005; Zheng et al., 2007), we wondered whether neural tuning for shape curvature, a property associated with mid-level visual area V4 (see Pasupathy et al., 2020), changed with age.

Visual areas V2 and V4 lie in the middle of the macaque form vision pathway. In adults, neurons in both areas are sensitive to stimuli containing conjunctions of orientations (Anzai et al., 2007; Gallant et al., 1993, 1996; Hegdé and Van Essen, 2000). More specifically, a number of studies have characterized the tuning of V4 neurons for curvature, with such neurons being sensitive to the curvature, either concave or convex, of segments within a larger bounded shape (Bushnell et al., 2011; Carlson et al., 2011; Namima et al., 2025; Nandy et al., 2013; Pasupathy and Connor, 1999). In V4, this tuning is often invariant to changes in scale (El-Shamayleh and Pasupathy, 2016) and position (Pasupathy and Connor, 1999). Such “object-centered” representations like these are thought to be critical for visual object recognition (see DiCarlo and Cox, 2007). By investigating how curvature is represented in the developing brain, including the extent to which curvature tuning is position-tolerant, we sought to improve our understanding of mature visual function.

To this end, we recorded multiunit activity from areas V2 and V4 of two developing macaque monkeys. We made longitudinal measurements during passive fixation at both 30 and 58 weeks of age while presenting shape stimuli that primarily varied in curvature at a single feature across multiple orientations and spatial positions (El-Shamayleh and Pasupathy, 2016). In addition to rate-based measures of curvature preference and position-invariant tuning, we modeled neural responses using both a deterministic, image-computable, linear model (akin to the spike-triggered average), and a stimulus-centric model of boundary curvature tuning, which has regularly been used to quantify the curvature tuning of V4 neurons. While we found curvature-driven responses in V2 starting at 30 weeks, these responses were spatially specific. On the other hand, we found evidence of object-centric curvature tuning in V4 from 30 weeks, the earliest age at which we made recordings. While both models explained a significant proportion of response variance at many sites across V2 and V4, model performance in both areas remained stable between 30 and 58 weeks. Our results suggest that V4’s object-centric tuning for curvature may be built from spatially-specific V2 afferents, all of which reach maturity relatively soon after birth.

## MATERIALS AND METHODS

### Data collection

We used data from two female *Macaca nemestrina* monkeys (M1 and M2). These animals were part of the same experimental series as Lee et al. (2024) and Rodríguez Deliz et al. (2025). Monkeys 1 and 2 here correspond to their monkeys 1 and 2.

Details of array placement can be found in Rodríguez Deliz et al. (2025) (see their Figure 2A, B). In short, we implanted 2 separate 96-electrode (Utah) arrays under sterile conditions. Electrodes were spaced 0.4 mm apart and were 1 mm in length. Following implantation, we used anatomical landmarks (including gross anatomical landmarks such as sulci and vasculature, which we observed surgically and related to previously documented area boundaries (Saleem and Logothetis, 2012; Winters et al., 1969)), physiological response properties (e.g. response latencies), and receptive field characteristics to determine the cortical location of array sites. In monkey M1, one array lay on the border between V1 and V2, the other in V4. In monkey M2, one array lay in V2, the other in V4. Because shape-evoked responses in V1 were typically unstructured, we did not consider them in this study. For completeness, we have included examples and summary data in Supplemental Figure 1.

**Figure 1:**
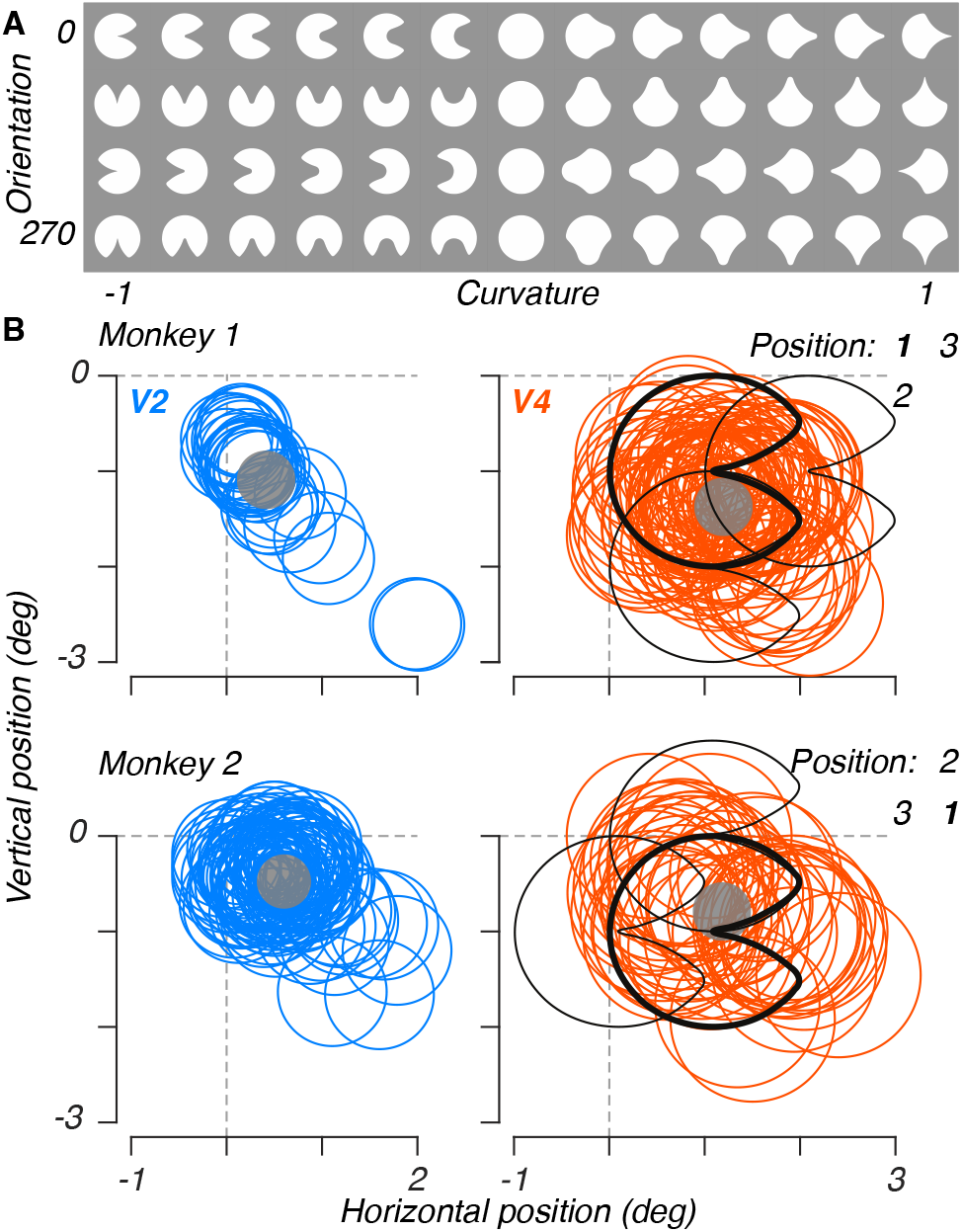
Stimuli and receptive field estimates. A: The 13 base shapes used in this experiment, which we presented at 4 orientations on a gray background. The curvature value associated with the principal feature is given below each unique stimulus. B: Receptive field estimates for both areas and animals (adapted from Rodríguez Deliz et al. (2025), Figure 2A, B). Each colored ring depicts the measured central position for each site. We measured a single common radius across all sites in a given area. Transparent gray circles indicate the average receptive field center for all sites in a given area. Black silhouettes indicate the three stimulus positions used for that animal. The bold silhouette shows the primary position used for all analyses; the other two were only used to measure position invariance.

**Figure 2:**
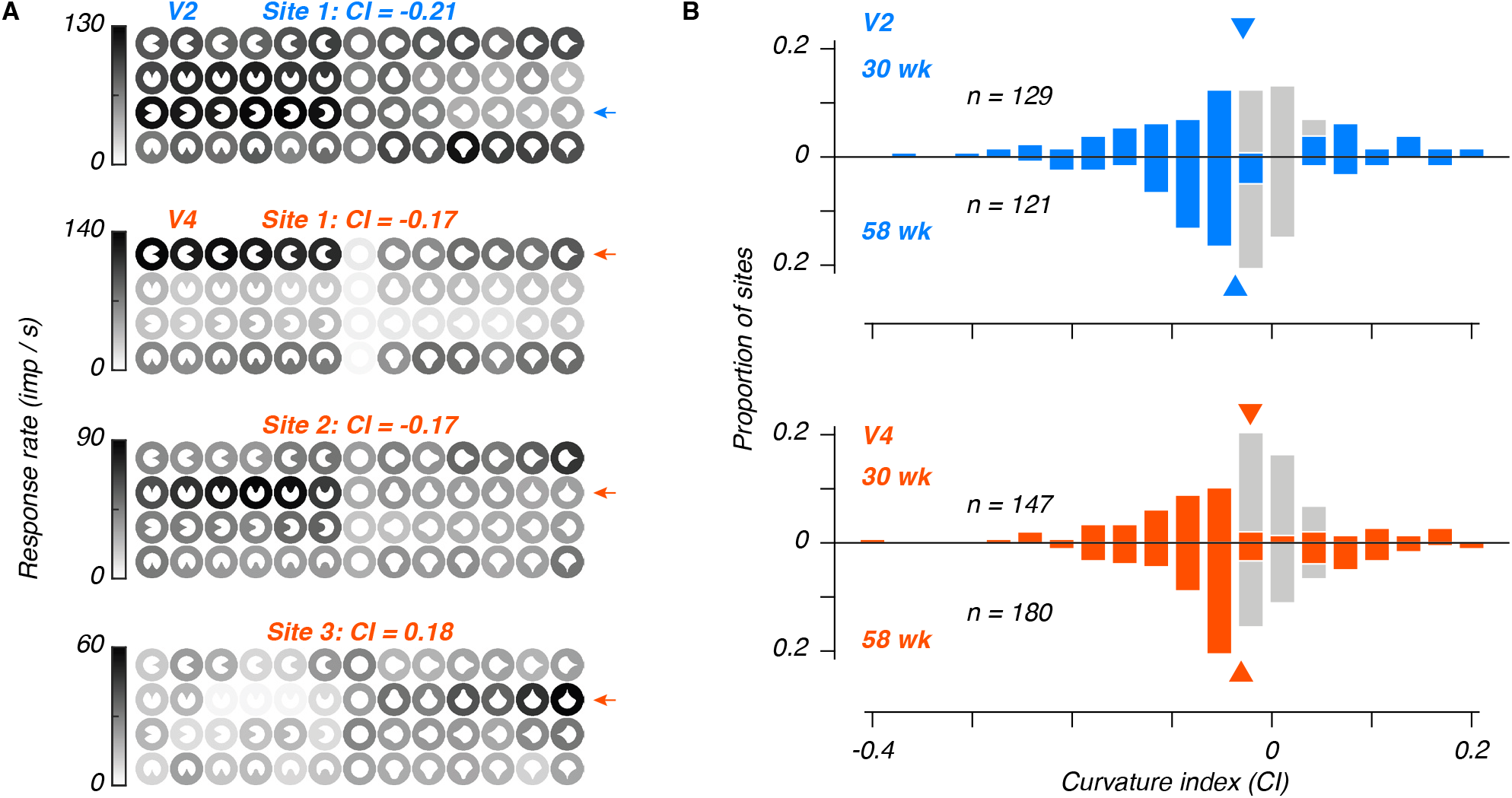
Curvature preference is stable between 30 and 58 weeks of age. A: Example responses measured at 30 weeks of age. Each shape’s background intensity represents the average response across repeated presentations of that stimulus. The arrow to the right of each panel depicts the preferred orientation for that site, which was used to compute the curvature index given at the top of each panel. B: Distributions of curvature indices across sites in V2 (top) and V4 (bottom). Data are plotted from 30 and 58 weeks (above and below the abscissa, respectively). Triangles depict median values. Colored bars depict sites whose responses are significantly stronger for either concave or convex stimuli, relative to shuffled null distributions.

We recorded neural activity from monkey 1 at 31 and 59 weeks of age, and monkey 2 at 29 and 57 weeks of age. We combined data from both animals at both ages. We recorded simultaneous multiunit spike events (threshold crossings) as voltage deviations exceeding 3 times the root-mean-square deviation of the baseline voltage. We performed all animal procedures in accordance with the National Institutes of Health *Guide for the Care and Use of Laboratory Animals* (2011), and with the approval of the New York University Animal Welfare Committee.

### Stimuli

We trained animals to passively fixate on a red square, 0.1 deg across. To first estimate receptive field locations (Figure 1B), we separately recorded neural responses to high contrast flashing dot stimuli, each one degree across, and presented across a 5 by 5 grid of locations, spaced one deg apart in both directions. We estimated the center location for each site, and fit a common radius across all sites from a given array. We used these receptive field estimates to determine where to position shape stimuli within the receptive fields of recording sites (below).

For our curvature experiments, we displayed the shape stimuli used by El-Shamayleh and Pasupathy (2016), which vary parametrically in boundary curvature (Figure 1A). We presented stimuli at a single spatial scale, corresponding to the ‘circle’ shape with a diameter of two degrees of visual angle, and four orientations, separated by 90 degrees and aligned with the primary curvature feature along a cardinal axis. Using our estimated receptive field locations, we displayed shapes at three positions – a central location, which we used for all analyses, and two secondary locations, which we used to estimate position invariance. The secondary positions were each one deg from the central location. In both animals, the central location was at (1°, −1°) relative to the center of gaze. In animal M1, the secondary locations were (1°, −2°), and (2°, −1°), as depicted in Figure 1B. In M2, the secondary locations were (0°, −1°), and (1°, 0°).

We presented stimuli 160 ms after fixation for 100 ms in duration, followed by a blank inter-stimulus interval of 100 ms. We presented stimuli in a pseudorandom order and sequentially in blocks of 8, resulting in a total block duration of 1760 ms. We presented all stimulus conditions (shape, orientation, and position) 30 times, with approximately 25% of stimuli being ‘blank’ images to estimate baseline firing rates.

We presented all stimuli on a gamma-corrected CRT monitor with a mean luminance of 28 cd m^-2^, a resolution of 1280 by 960 pixels, and a frame rate of 100 Hz. We seated animals in a custom primate chair 114 cm from the monitor; the monitor subtended 20 by 15 deg.

### Analysis

#### Visually-responsive sites

For all analyses, we summed the spike count from 50 to 200 ms following stimulus onset; this window covered the time from the beginning of the neural response to the beginning of the following stimulus presentation. We subtracted the baseline response, measured using responses to blank stimuli, for all analyses. To determine whether sites were visually responsive, we used both a measure of responsiveness from baseline (*d*′) and a measure of tuning reliability (split-half Pearson correlation). First, we measured the difference between the response distributions for blanks and all stimuli presented at the central position, as

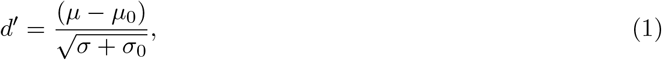

where *µ, σ* and *µ*_0_, *σ*_0_ are the mean and standard deviation of responses to stimuli and blanks, respectively. To measure split-half correlation for each site, we extracted response vectors for all stimuli at the central position (excluding blanks), constructed from the average of even and odd stimulus repetitions. Finally, we compute the Pearson correlation between these two response vectors to quantify response reliability.

For both metrics, we used permutation tests to assess significance (two-tailed for *d*′, one-tailed for split-half correlation; *α* was 0.05 for both), and included all sites meeting at least one of these significance criteria for further analysis.

#### Curvature index

As a model-free metric of curvature tuning, we computed a Curvature Index (CI) for each site based on the response difference between concave and convex stimuli. Specifically, at the orientation eliciting the highest mean response, we compute the average activity for stimuli with similarly signed curvature, denoted *r*^+^ (convex) and *r*^−^ (concave), *i*.*e*.

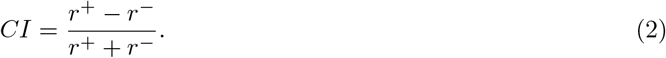

Curvature indices were bounded by [-1, 1], where negative and positive values indicate concave- and convex-preferring sites, respectively. We estimated confidence intervals using a bootstrap procedure, computed across 10000 resamples.

#### Position invariance

To understand how stimulus shifts in visual space affected neural response characteristics, we computed the Pearson correlation for the response vectors to all stimuli at each pair of visually-responsive positions for a given site (including the two secondary locations). For sites responsive to all 3 locations, we analyzed each pair independently. To determine significance, we permuted the response vector across shape stimuli for each site and position 10,000 times, and used each pair of permuted response vectors to compute a null distribution of correlations. We considered any positional correlation outside the 95th percentile of this null distribution to be significant.

#### Angular Position and Curvature (APC) model

We fit the angular position and curvature (APC) preference of each visually responsive site using the model of Pasupathy and Connor (2001). To map stimuli into the APC space, we discretize the 8 diagnostic features of a stimulus, located approximately 45 degrees apart along the boundary, half of which are aligned to the cardinal axes (see El-Shamayleh and Pasupathy, 2016). To calculate curvature at these locations, we used a piecewise linear approximation of the boundary turning angle, followed by a compressive nonlinearity to map these values onto the curvature interval between −1 (concave) and +1 (convex):

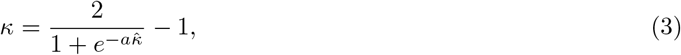

*κ* and 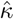 represent the compressed curvature and raw turning angle, respectively. The *compression coefficient a* was defined as *a* = 0.0483 following Pasupathy and Connor (2001). This process yielded 8 coordinate pairs of angular position and curvature for each stimulus, spaced (approximately) evenly in polar angle and aligned with the cardinal axes. Specifically, for a stimulus Γ, we approximate its corresponding shape contour with Γ′, a vector of points (*γ*_*θ*_, *γ*_*κ*_) in the space of angular position and curvature.

The APC model maps neuronal shape tuning using a 2-dimensional separable Gaussian function in the angular position and curvature space (Oleskiw et al., 2018, 2014; Pasupathy and Connor, 1999). The Gaussian function is circular in the domain of angular position. Here, the predicted response to a stimulus is determined by evaluating the maximum activation of each contour point:

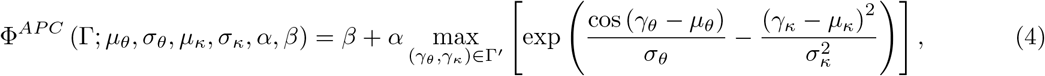

with *µ*_*θ*_, *µ*_*κ*_, *σ*_*θ*_, and *σ*_*κ*_ being the centers and spreads of the angular position and curvature tuning function. To match the range of neural responses observed in the data, we also include gain (*α*) and offset (*β*) parameters.

#### Linear receptive field (LRF) model

For comparison, we also consider a baseline image-computable deterministic model to relate neural response *r*_*i*_ to the particular luminance patterns of the *i*^*th*^ stimulus image Γ_*i*_. Simply put, we measure the spike-triggered average *ψ* across all stimuli as

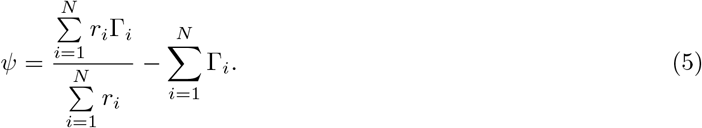

Here, the template *ψ* estimates a receptive field from the set of *N* stimuli that is linear in pixel space. With this template, a neuron’s response to a stimulus Γ is predicted as simply:

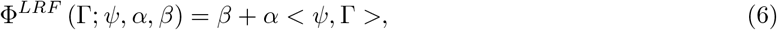

*i*.*e*., the inner product between the template and stimulus. To map the arbitrary output units of the two models to a response rate, we again parameterize a gain *α* and offset *β*.

Note that we systematically downsampled the stimulus images and fitted the LRF model to assess the impact of the number of template coefficients on model performance. We observed that the model’s performance plateaued at a 10×10-pixel stimulus representation and used this resolution for the LRF model analysis.

#### Model optimization and evaluation

For each neuron, we optimize the parameters of our two models to minimize the error between the average evoked response and model prediction across our stimuli. Thus, for our stimulus set {Γ_*i*_}|*i* ∈ 1, …, *N* and corresponding response to the *i*^*th*^ stimulus *r*_*i*_ we seek the set of optimal model parameters

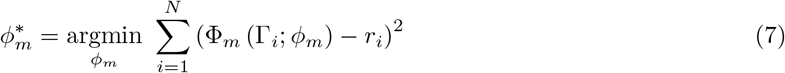

for each model *m*, either APC or LRF. Both sets of optimal model parameters are estimated simultaneously using a simplex-based deterministic gradient descent. Note that the APC model has six independent parameters, while the LRF model has only two. To avoid local minima solutions, the optimization procedure is repeated with initial conditions sampled uniformly over the parameter space (Oleskiw et al., 2014).

To select among candidate models, we used a 13-fold cross-validation to assess how well each model generalizes to novel stimuli. Each training set contained 48 of 52 possible stimuli (including all shapes and orientations), randomly selected. We used the 4 held-out stimuli for testing, with each shape held out exactly once. We measured model performance as the explained variance (*r*^2^) averaged across test sets. To then relate performance between the two models, we calculated the partial correlation (El-Shamayleh and Pasupathy, 2016; Movshon et al., 1985; Smith et al., 2005) with:

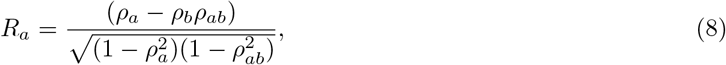

where *a* and *b* denote either model. Specifically, *ρ*_*a*_, *ρ*_*b*_ is the Pearson correlation between observed responses and predictions of model *a,b*, and *ρ*_*ab*_ is the correlation between the predictions of both model *a* and *b*.

To measure the significance of each partial correlation *R*, we utilize the Fisher *z*-transformation, *e*.*g*.,

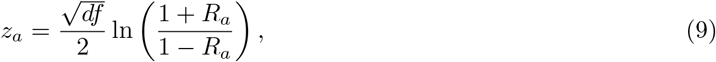

which is computed for each model. Note that to standardize *z*, it is scaled by the inverse square root of the degrees of freedom of the correlation calculation, determined by the number of stimuli minus 3, *i*.*e*., 49. If, for a given site, the difference between *z* values was at least 1.28, equivalent to exceeding the 90^*th*^ percentile of randomly sampled correlations, the site was deemed to be a better fit by the model with a greater *z*. If both *z* values were below 1.28, neither model was significantly superior.

## RESULTS

To assess whether neural tuning for boundary curvature changed with development, we recorded multiunit neural responses in V2 and V4 to shape stimuli whose curvature differed primarily along a single boundary segment. We collected longitudinal data from 2 macaque monkeys at 30 and 58 weeks of age. To quantify the boundary shape of these stimuli, we measured curvature along these boundary segments and normalized values to range from concave (-1) to convex (+1). We presented each stimulus at four orientations during passive fixation (Figure 2A), and measured neural tuning for curvature. We also quantified the stability of tuning across positional shifts, and present two models to explain the data.

### Curvature tuning is diverse across sites, but stable across ages

Figure 2A shows responses from 4 example sites (1 in V2, 3 in V4) to all stimuli. Tuning across both curvature and orientation varied across sites: some sites responded to a range of curvatures and orientations (*e*.*g*., V2 site 1, V4 site 1), while others exhibited sparse activations (V4 sites 2 and 3). In V4, responses were often highly structured within the 2-dimensional stimulus space, with well-defined maxima (V4 sites 1, 2, and 3). In V2, responses were typically less structured (V2 site 1), though this was not always the case (V2 sites 2, 3, and Supplemental Figure 2A). Finally, as stated above, one V2 array was situated on the V1/V2 border. Supplemental Figure 1A shows two example sites from V1. Because sites in V1 were, as expected, unstructured in their responses to curvature stimuli, their analysis was included as supplementary material.

We measured curvature tuning of each site in V2 and V4 using a curvature index (the difference between convex and concave responses, normalized to their sum). We measured this index at the orientation that elicited the highest average response, yielding values ranging from -1 (concave-preferring) to +1 (convex-preferring). Note that this value reflects the preference of a site for convex versus concave curvature, but does not reflect the sharpness of a site’s tuning within those two categories. The curvature indices we measured in Figure 2A were similar in magnitude, even as they varied in the width of their selectivity within their preferred curvature sign. We observed a wide range of curvature indices across our sample (Figure 2B) for both V2 and V4 at 30 and 58 weeks. These ranges remained stable across age, as did the medians of the distributions (median, [95% CI]: *V* 2_30*wk*_ = −0.03 [−0.04, −0.01]; *V* 2_58*wk*_ = −0.04 [−0.05, −0.03]; *V* 4_30*wk*_ = −0.02 [−0.03, −0.01]; *V* 4_58*wk*_ = −0.03 [−0.04, −0.02]). We found no evidence that curvature tuning became stronger after 30 weeks of age.

### Position-invariant tuning in V4

Neural tuning for curvature in V4 is often invariant to positional shifts (within the receptive field, see Pasupathy and Connor (1999)). We wondered whether this position invariance in V4 might change with age, and to what extent the curvature tuning we observed in V2 was position-invariant. We presented stimuli at three total positions: the primary position used above, and two others, each offset from the primary position by 1 deg of visual angle (approximately 50% of the total stimulus size; see Figure 1B). For any site that was responsive to stimuli at more than one location, we computed the correlation between responses at the response-evoking locations (including the two secondary positions, if all positions evoked a response).

Responses corresponding to all three positions are shown in 3A for example sites within V2 and V4. Note that while this V2 site responded strongly to all three positions, the response patterns were largely uncorrelated across shapes. The examples shown in Supplemental Figure 2A also varied widely across position.

Figure 3B shows all the positional correlations we measured across our entire sample of V2 and V4 sites. As in our examples, curvature tuning in V2 was rarely stable across positional shifts: the median positional correlation was close to 0 at both ages (-0.00 95% *CI*_30*wk*_: [-0.02, 0.02]; *CI*_58*wk*_: [-0.04, 0.02]). For each pairwise comparison, we performed a permutation test on the response vectors to assess whether the correlation was significant, compared to an *α* of 0.05. In V2 at 30 weeks, 9% (95% CI: [6.19%, 12.69%]) of pairwise comparisons were significantly positively correlated; 7% (95% CI: [4.33%, 10.21%]) were negatively correlated. At 58 weeks, 6% (95% CI: [3.50%, 9.34%]) were positively correlated; 9% (95% CI: [5.83%, 13.23%]) were negatively correlated. At both ages, the number of positively correlated comparisons was close to our choice of *α*, and was (approximately) the amount expected by chance.

**Figure 3:**
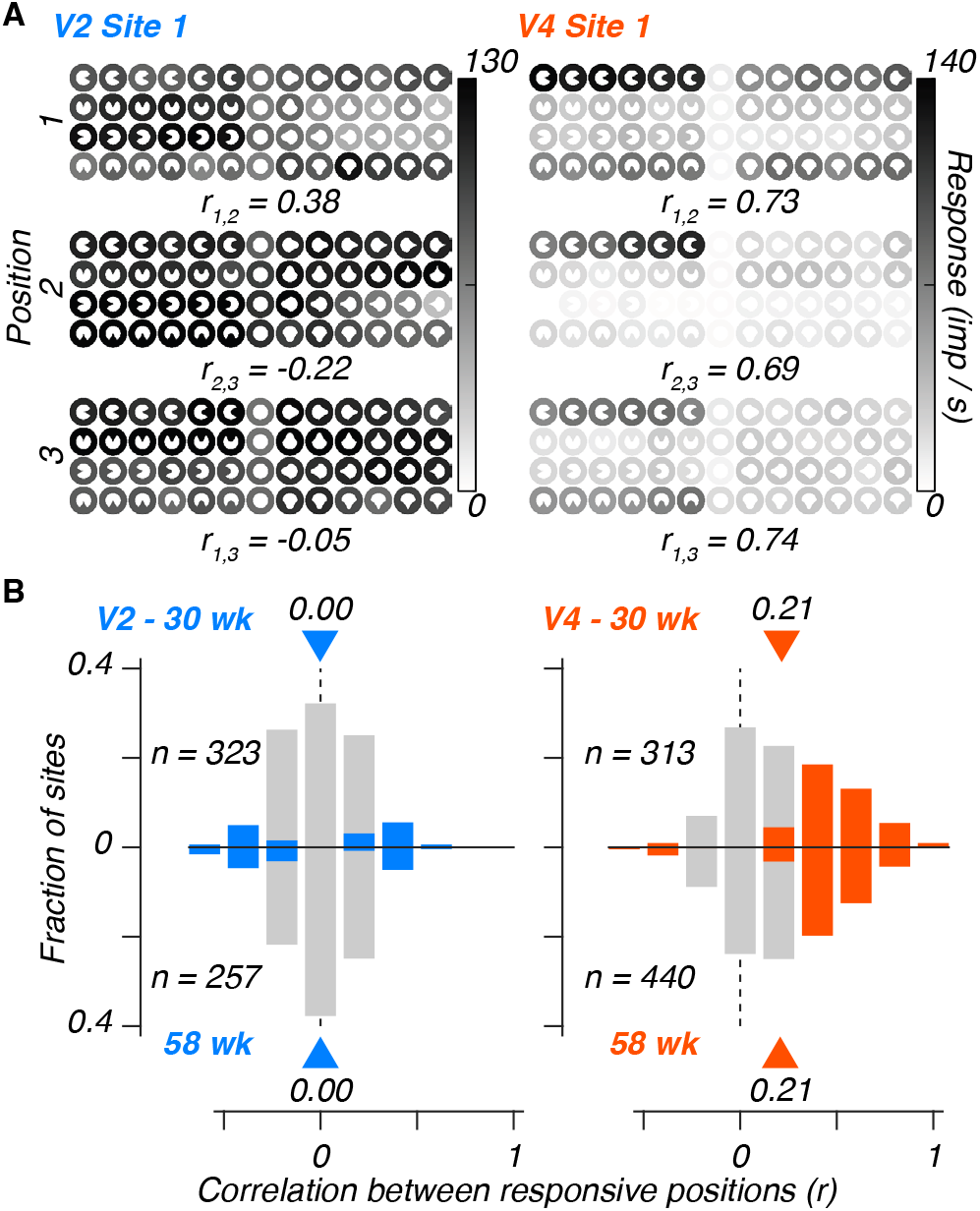
Position invariance in V4. A: Average responses for example V2 (left) and V4 (right) sites at three positions, measured at 30 weeks of age, depicted as in Figure 2A, with the three positions shown in Figure 1B. Numerical values between panels are response correlations between positions. B: Distributions of position correlations for both ages measured, with histograms above the abscissa depicting correlations from sites measured at 30 wk, and below depicting the same measured at 58 wk. Triangles and associated numerical values indicate the median correlation for that sample. Colored bars represent sites with significant correlations between visually-responsive positions, and gray bars represent the proportion of sites without significant correlations.

In V4, the median correlation was positive, and was identical at both ages (0.21 in both cases; 95% *CI*_30*wk*_: [0.17, 0.26]; *CI*_58*wk*_: [0.16, 0.23]). At 30 weeks, 42% (95% CI: [37.06%, 47.92%]) of positional comparisons were positively correlated; 1% (95% CI: [0%, 2.23%]) were negatively correlated. At 58 weeks, 40% (95% CI: [35.23%, 44.32%]) of positional comparisons were positively correlated, while 3% (95% CI: [1.14%, 4.09%]) were negatively correlated. Unlike in V2, we observed several sites in V4 that were tolerant to shifts in stimulus position. Thus, position invariance was stable from 30 weeks of age in V4, and was not observed in V2.

### Modeling shape selectivity in V2 and V4

Using direct measurements of neural responses (*i*.*e*., CI and position invariance), we found evidence that curvature tuning was stable from 30 weeks of age. Next, we wondered whether models of neural curvature tuning might reveal quantitative signs of development. We fit two models: the first is the Angular Position and Curvature (APC) model (Pasupathy and Connor, 2001), which predicts neural responses using tuning functions over a two-dimensional representation of a stimulus’s curvature along its circumference. The second Linear Receptive Field (LRF) model is an image-computable model based on the spike-triggered average response to luminance across the stimulus. We fit both models to our recordings from V2 and V4, at 30 and 58 weeks. We used non-overlapping training and testing subsets of our stimulus set to measure the cross-validated performance of each model.

### Angular position and curvature (APC) model

The APC model of Pasupathy and Connor (2001) assumes separable tuning for the curvature of a contour segment (concave or convex), and the relative orientation (angular position) of that segment relative to the rest of the contour. The tuning of a given site is captured by the product of two separable Gaussian tuning functions. An additional gain and offset parameter maps the model output to the neural response range.

Figure 4A represents the model fit, within the APC space, for the example sites shown above. As an example, for V4 site 1 (second column), the model prediction corresponds to a site broadly responsive to concave boundary segments (negative curvature values) centered between 270 and 0 degrees. Figure 4B represents the predicted responses to all stimuli. Model performance, measured as the cross-validated variance explained (*r*^2^), was high (0.85). APC model performance varied across sites. While V4 site 2 (Figure 4A, third column) was also well described by the model, the fit for V4 site 3 (Figure 4A, right column) captured the sparseness, but not the peak, of the recorded site.

**Figure 4:**
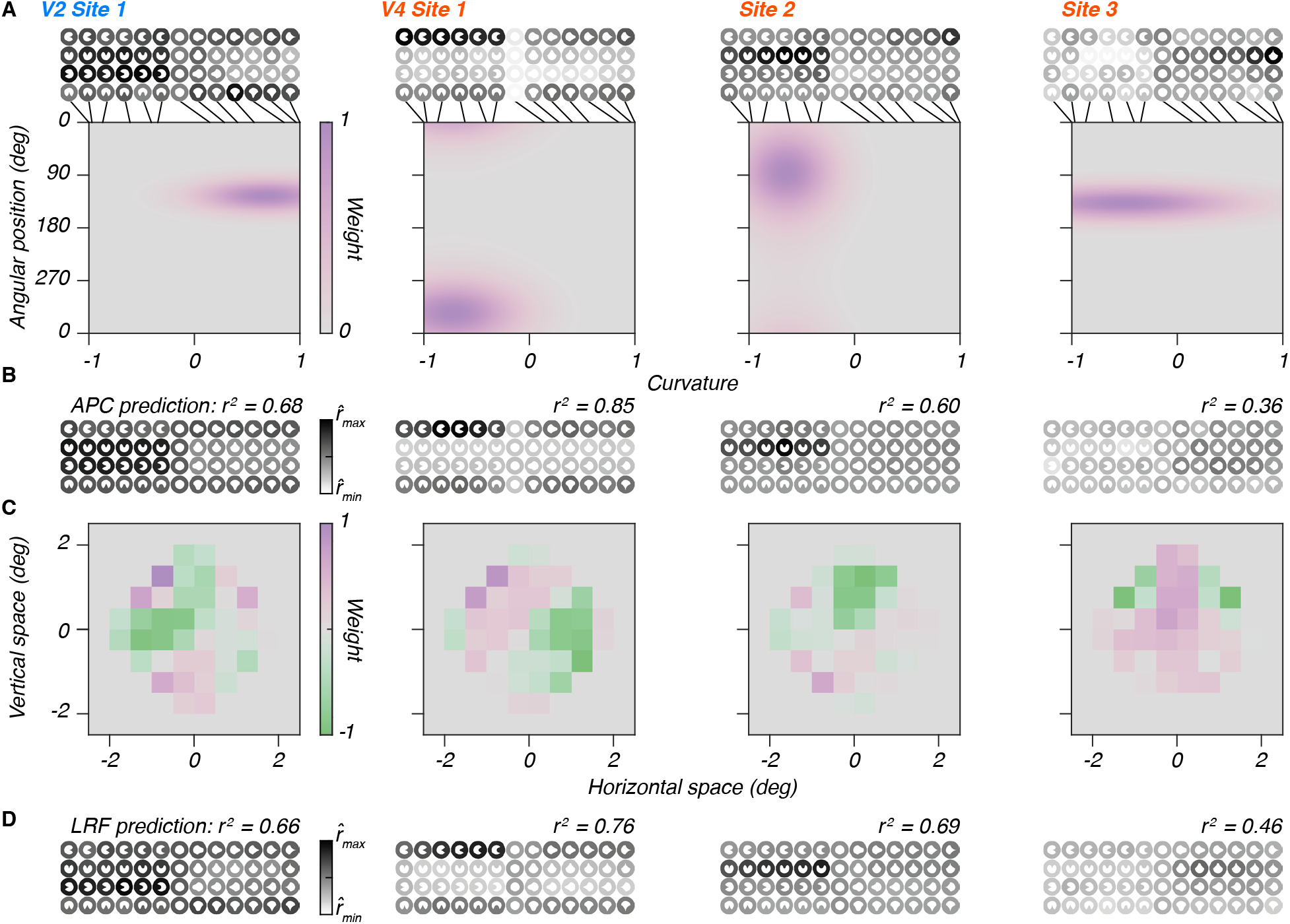
Example fits for V2 and V4. A: Example APC model fits for sites in V2 and V4, fit to all data. Shading represents the relative weight for each location in the space of angular position and curvature. Example sites are the same as in Fig. 2A; inset panels (top) show the actual responses of each site. The ticks at the top of each panel mark the measured curvature values for the 13 shapes. B: APC model predictions for the example sites shown in Figure 2A, with cross-validated variance explained. C: Example LRF model fits, based on all stimuli (not cross-validated). The color saturation of each pixel represents its relative weight. D: Cross-validated LRF model predictions for the same example sites.

We also evaluated the APC model’s performance in V2. The model fit for V2 site 1 (Figure 4A, left column) predicts a tuning for convex boundary segments between 90 and 180 deg. Our tiling of angular position was too sparse to measure responses to convex stimuli at this position. On the other hand, our *concave* stimuli, when presented at either 90 or 180 deg, have a convex feature at that angular position. Hence, the model paradoxically predicts that concave, not convex, stimuli will elicit the strongest responses at the site. This prediction comports with our measurements for the site (Figure 4B, left column), but also demonstrates that our sampling of angular position was too sparse to evaluate APC predictions fully. The accurate predictions in V2 sites 2 and 3 (Supplemental Figure 2B) are also subject to this caveat. In V1 (Supplemental Figure 1B-C), the APC model made predictions which were inconsistent with the boundary curvature tuning we observed in V4, regardless of overall model performance: V1 site 1 was the V1 site best predicted by the model (*r*^2^ = 0.87), but the predicted response pattern was broad and included concave and convex sites at multiple orientations, as opposed to an orderly response to a limited set of orientations and curvatures *e*.*g*. V4 sites 1-3, V2 sites 1-3 (Supplemental Figure 2A).

### Linear receptive field (LRF) model

We found examples of sites in all areas whose responses could be well described in the APC model space of stimulus-centric boundary curvature. To compare the APC model’s performance, we developed a simple image-computable model based on the spike-triggered average response to each pixel. This Linear Receptive Field (LRF) model can be thought of as a simplification of other linear and linear-nonlinear models, which have previously been used to model curvature tuning in V4 (Cadieu et al., 2007). For this model, we downsampled input images and measured the spike-triggered average weight for each pixel in the stimulus domain. This generated a filter containing positively (on) and negatively (off) tuned subregions. The inner product of this filter with a given stimulus yields the predicted response to that stimulus. We then added gain and offset parameters to map filter outputs to the neural response range.

Figure 4C represents the normalized filters for each receptive field subregion, for the same 4 example sites depicted previously. Starting with V4 site 1 (second panel), we extracted a filter corresponding to excitatory and inhibitory lobes approximately 135 deg and 0 deg from horizontal, respectively. The LRF model therefore predicts strong responses to concave features at 0 deg, as represented in Figure 4D, and captures 76% of the overall response variance to our stimuli. The model prediction for V4 site 2 is of strong responses to concave boundary segments at 90 deg, matching our measurements. The model prediction for V4 site 3 includes positive subregions, flanked in part by negative lobes above the horizontal midline. The prediction is of a maximum response for stimuli with convex boundary segments positioned at 90 deg, but, as with the APC prediction, the model fails to fully predict the responses to all stimuli in the set.

For V2 site 1, we extracted a filter with inhibitory lobes at the top and left sides, and excitatory lobes in the upper-left, upper-right, and lower-left corners. The resultant model prediction matched the data, except for the single convex stimulus (at 270 deg), which elicited a strong response not captured by either model. V2 sites 2 and 3 (Supplemental Figure 2C) were also relatively well described by the model. In V1, we extracted filters with adjacent, parallel lobes with opposing tuning (Supplemental Figure 1D-E), which resembled standard models of orientation tuning in V1 (DeAngelis et al., 1993). These examples demonstrated that the models could capture a significant fraction of the response variance, even if some model predictions deviated from the data (*e*.*g*. the sparseness of V4 site 3, the convex response in the V2 site).

### Model performance remains stable from 30 weeks

We fit both models to all visually responsive sites in V2 and V4. Figure 5A shows the variance explained (*r*^2^) by each model. We next used Fisher’s r-to-Z transformation to convert each to units of standard deviation (as in El-Shamayleh and Pasupathy, 2016; Smith et al., 2005). This conversion allowed us to relate differences in partial correlations, independent of their overall magnitudes, to isolate the amount of explainable variance uniquely captured by one model or the other. Furthermore, we may determine which model, if any, captured a significant portion of response variance. We used a threshold of *α* = 0.1 to assess significance.

**Figure 5:**
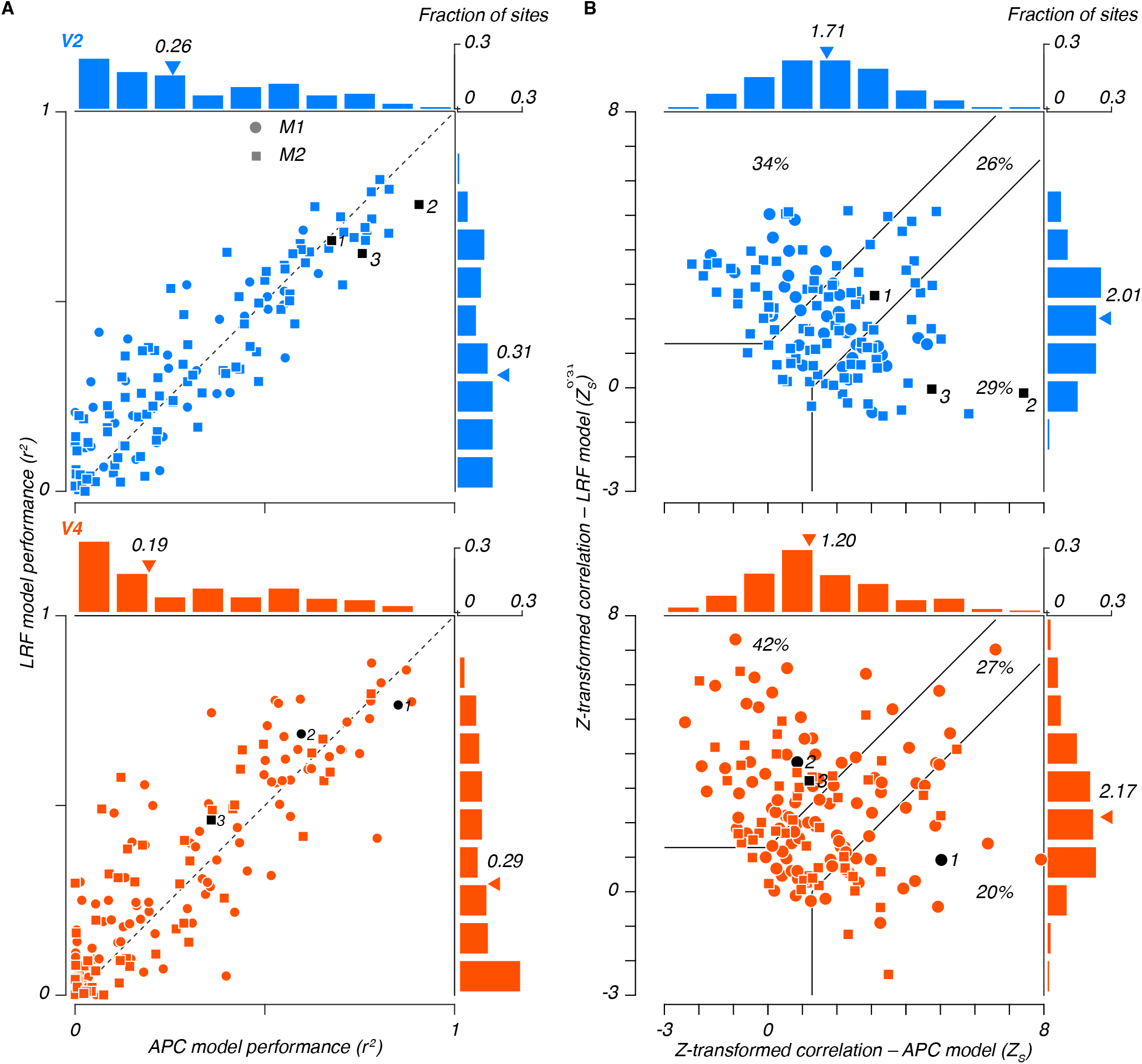
Model performance and comparison. A: Cross-validated variance explained for each model, for each site in V2 (top) and V4 (bottom), for data collected at 30 weeks from M1 (circles) and M2 (squares). APC model values are plotted along the abscissa; LRF values along the ordinate. Marginal histograms show the distributions of these values; triangles indicate their medians. The example sites from prior figures are numbered and plotted in black. B: Z-transformed partial correlations from each model. Significance bounds (black lines) denote sites explained well by either model.

Partial correlations are shown in Figure 5B, for data collected at 30 weeks. Based on our threshold, we categorized all sites into one of four possible categories: *i)* better explained by the LRF model (above the upper boundary), *ii)* better explained by the APC model (below the lower boundary), *iii)* well explained by both (between the slanted segments of the bounds), and *iv)* not explained by either model (exceeding neither vertical nor horizontal bound components).

Across our samples in both V2 and V4, the majority of sites (89% in V2, 89% in V4) were well captured by at least one model (5B, sites beyond either significance bound). In V2 (5B, top), a plurality of sites were well captured by both models (between bounds, 26%); similar quantities of sites were uniquely described by either model independently (upper-left LRF: 34%, lower-right APC: 39%). In V4 (5B, bottom), many sites were well captured by both models (between bounds, 27%). A plurality of sites were well captured by the LRF model (upper-left, 42%), and a smaller fraction was uniquely captured by the APC model (lower right, 20%). In V2, the median z-transformed partial correlations were (5B, marginal distributions) *Z*_*APC*_ = 1.71 (95% CI: 1.30-2.10) and *Z*_*LRF*_ = 2.01 (95% CI: 1.63-2.42), and in V4 were *Z*_*APC*_ = 1.2 (95% CI: 0.85-1.48) and *Z*_*LRF*_ = 2.17 (95% CI: 1.76-2.73). In contrast, the overall performance of both models for sites in V1 was relatively low, yet 84% of sites were well-fit (Supplemental Figure 1F-G) by at least one model.

We again fit our models to data collected at 58 weeks, determining the fraction of sites which could significantly be explained by either model type (Supplemental Figure 3: sites which were captured by both models are counted in both fractions). As with curvature index and position invariance, we found no evidence of age-related changes in the fractions of sites explained by either model in either area.

## DISCUSSION

In these experiments, we found that neural representations of boundary curvature in macaque V2 and V4 are adult-like from 30 weeks of age, the earliest ages we tested. We also found that tuning for curvature in V4 was invariant to stimulus position; this tuning was also adult-like by 30 weeks of age. In V2, responses were spatially specific at all ages. We modeled responses using both descriptive (APC) and image-computable (LRF) models. We found no evidence that model performance changed as a function of age, nor did the relative performance of each model change with age.

### Visual development

Measurements taken in the LGN and V1 of developing macaques suggest that tuning properties mature by or before 16 weeks (Kiorpes and Movshon, 2004; Maruko et al., 2008; Movshon et al., 2005). In downstream V2, data suggest that development may be slower than in V1 (Maruko et al., 2008), but that tuning properties quickly reach maturity (Kiorpes and Movshon, 2004; Lee et al., 2024; Maruko et al., 2008; Rodríguez Deliz et al., 2025; Wang et al., 2019; Zhang et al., 2013, 2005; Zheng et al., 2007). Our results in V2 add further support to the notion that neurons in area V2 mature soon after birth.

How neuronal tuning in area V4 develops has remained largely unexplored – we previously reported that selectivity for naturalistic texture statistics (Lee et al., 2024) and for radially modulated shape patterns (Rodríguez Deliz et al., 2025) is present from 30 weeks. The results reported here, taken from the same animals, expand this notion of rapid maturity to include curvature tuning, and position invariance.

Analysis of the response dynamics of V4 neurons suggests that tuning for the types of curvatures used here may arise through recurrent processing, either in V4, or a combination of V4 and posterior inferotemporal cortex (pIT) (Yau et al., 2013). We have previously reported that the sensitivities of V4 and pIT neurons to naturalistic texture statistics are mature by 30 weeks. This sensitivity is apparently extracted in a feedforward manner (Lee et al., 2024; Ziemba et al., 2019). If indeed curvature tuning is based on recurrent processing in or near V4, then our results suggest that these mechanisms may also emerge soon after birth.

#### Curvature tuning in V2 and V4

Previous studies have demonstrated that neurons in V2 can be tuned for complex shapes (Hegdé and Van Essen, 2000), and conjunctions of angles (Anzai et al., 2007; Ito and Komatsu, 2004). Here, we found that neurons in V2 are also sensitive to the curvature of shape boundaries, but in a manner which is location-specific. These results provide a natural platform on which object-centric curvature tuning, often observed in V4 (Cadieu et al., 2007; David et al., 2006; Gallant et al., 1993; Pasupathy and Connor, 1999; Sharpee et al., 2013), may be built. Our interest was the study of development; thus, we were unable to probe this connection further (experimental constraints prevented us from tailoring our stimuli more closely to the size and position of our V2 receptive fields). Recent advances in both recording techniques (Kim et al., 2026; Namima et al., 2025), and closed-loop stimulus generation (Cowley et al., 2026) have led to a deeper understanding of the organization and function of both V2 and V4. Moving forward, these advances may support future efforts to better illuminate the origin and mechanisms supporting object-centric tuning in the midlevel visual cortex.

## Acknowledgements

We are grateful to members of the Visual Neuroscience Laboratory for advice and discussion. This work was supported by grants from the National Institutes of Health: R01EY022428 (J.A.M.), R01EY024914 (L.K. and J.A.M.), R01EY031446 (N.J.M.), F31EY031249 (G.M.L.), as well as Fellowships from the NSF-REU (under award number 2447707), the Simons Collaboration on the Global Brain, and the Flatiron Institute (to A.E.S.).

## Author contributions

N.J.M., L.K., and J.A.M. designed research; G.M.L. performed research; A.E.S. analyzed and modeled the data, guided by G.M.L., T.D.O., and N.J.M; A.E.S. and G.M.L. drafted the paper; and T.D.O., J.A.M. and L.K. finalized the paper.

## SUPPLEMENTARY FIGURES

**Supplementary Figure 1:**
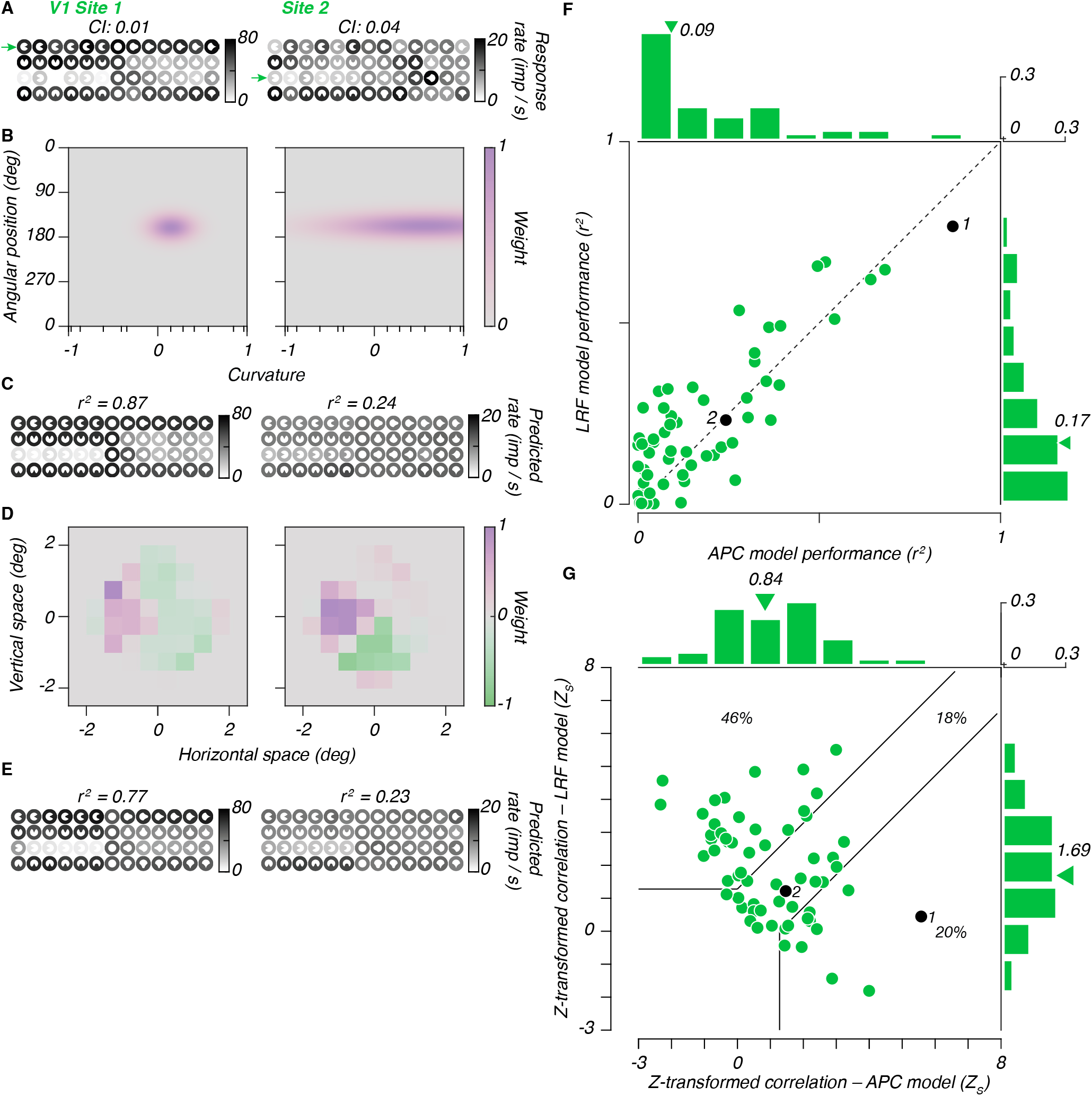
Examples and summary analyses based on recordings taken in V1. A: Example responses measured at 30 weeks of age (conventions as in Fig. 2A). B: APC model fits for example sites in A, fit to all data (not cross-validated). Conventions as in Fig. 4A. C: Cross-validation APC model predictions for the example sites shown in B. Conventions as in Fig. 4B. D: Example LRF model fits, based on all stimuli (not cross-validated). Conventions as in Fig. 4C. E: Cross-validated LRF model predictions for the example sites shown in D. Conventions as in Fig. 4D. F: Variances explained for both models, for each visually responsive site in V1, for data collected at 30 weeks. Conventions as in Fig. 5A. G: Z-transformed partial correlations for both models. Conventions as in Fig. 5B.

**Supplementary Figure 2:**
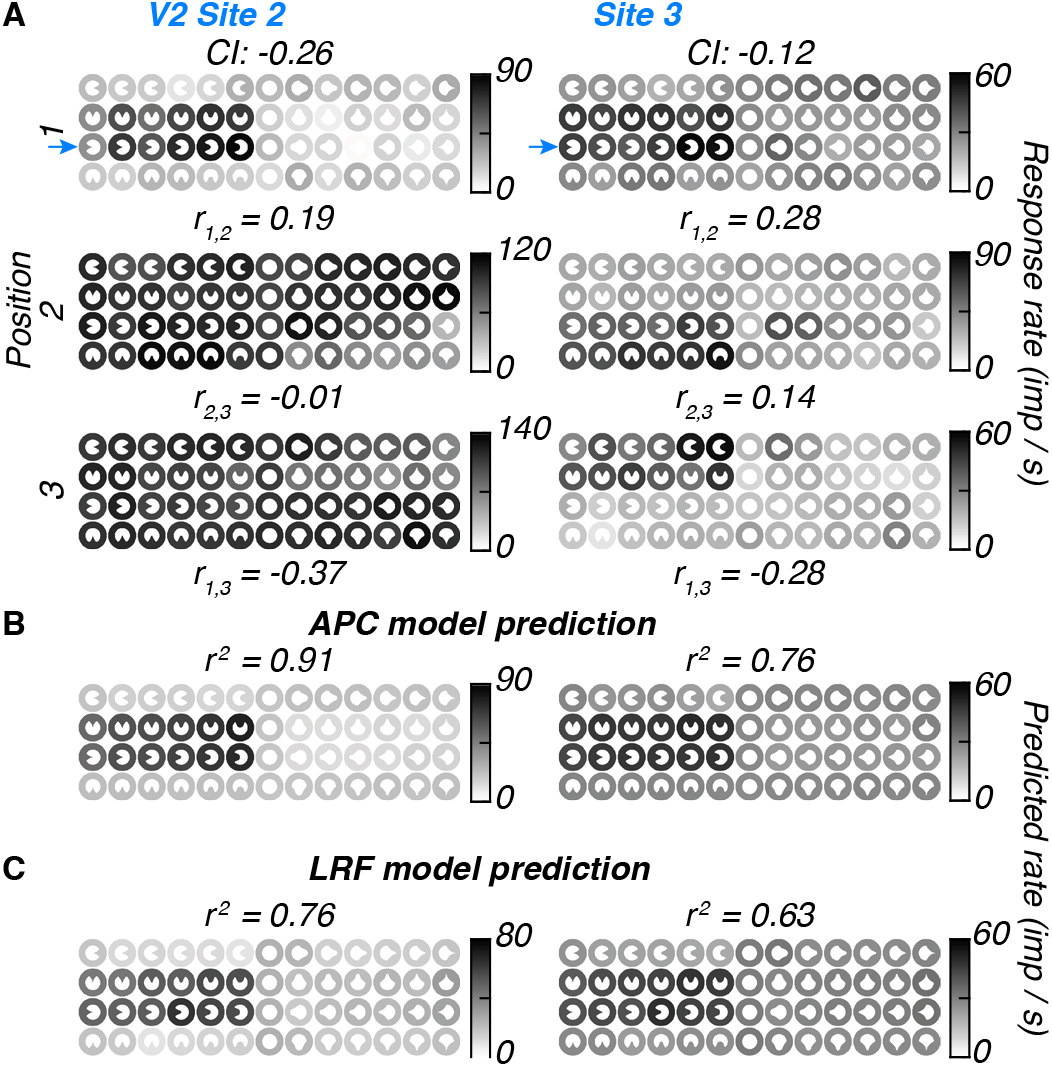
Additional representative sites recorded from V2. A: Responses to the three presentation locations. Conventions as in Fig. 3A, though note that the different stimulus positions are scaled to different ranges. B: APC model predictions for the example sites in A, for the primary stimulus position. Conventions as in Fig. 4B. C: LRF model predictions for the example sites in A, for the primary stimulus position. Conventions as in Fig. 4D.

**Supplementary Figure 3:**
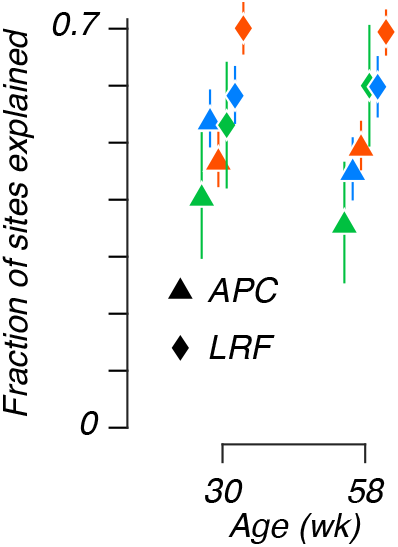
Fraction of sites explained by the models, grouped by age of recording. Data points were horizontally shifted for visibility. Sites well-captured by both models were counted twice.

